# How air temperature and solar radiation impact life history traits in a wild insect

**DOI:** 10.1101/2024.08.13.607703

**Authors:** Alexandra S. Gardner, Ilya M.D. Maclean, Rolando Rodríguez-Muñoz, Alfredo F. Ojanguren, Tom Tregenza

## Abstract

Ectotherms are essential components of all ecosystems and rely on external heat to regulate their body temperature. For most terrestrial ectotherms the primary sources of heat are ambient temperature and solar radiation. Many insects can use movement to respond to changes in temperature and solar radiation in order to manage their body temperature and optimise life history traits. However, we lack the understanding of the relative importance of temperature and shade that we need to predict how the combined effects of changes in air temperature and cloud cover will impact terrestrial insect populations. We reared developing nymphs of the field cricket (*Gryllus campestris*) at high and low air temperature sites with partially shaded and unshaded treatments at each site. Given the broad altitudinal range of this species, we tested the possibility of local adaptation to these climate variables by rearing nymphs from high and low altitude genetic lineages in all treatment combinations. We found that development time was strongly affected by air temperature, but not by a substantial increase in shade. This suggests that developing crickets can compensate for an increase in shade, presumably because in unshaded conditions they forgo some opportunities to gain energy from the sun. We found that mass at adulthood was affected by an interaction between availability of sun (shading treatment) and air temperature. This indicates that changes in cloud cover will impact insects differently in warmer and cooler areas. We found no evidence for local adaptation in these traits. Our findings underscore the need to consider both ambient temperature and solar radiation in predicting the impacts of climate change on insect populations, as shifts in temperature and cloud cover may have complex and region-specific effects on these vital ecosystem components.

## Introduction

The thermal environment profoundly impacts ectotherms because they rely on external sources of heat to regulate their body temperature (Angiletta, 2009). Air temperature plays a crucial role in determining the availability of suitable thermal environments. Solar radiation is also important because basking in sunshine can raise body temperature by more than 20°C above ambient air temperatures (Gardner et al., 2024). Many ectotherms bask to maximise radiative heat gain (Lahondère, 2023).

The combination of air temperature and solar radiation has been observed to explain the distribution and abundance of insects at various spatial scales (Bryant & Shreeve, 2002; Kührt et al., 2006). However, few studies have investigated how they interact to affect life history and fitness related traits, which ultimately drive these patterns (e.g., thermal stress: Buckley et al., 2013). It is important to bridge this knowledge gap because ectotherms make up the majority of terrestrial biodiversity (Wilson, 1992) and so any changes to the thermal environment will have implications for biological systems globally.

Temperature and solar radiation are not identical in their potential to affect fitness. Solar radiation often varies enormously at small spatial scales through availability of shade. Sun basking is a critical part of thermoregulation for many insect species, enabling them to reach optimal body temperature for physiological processes such as digestion, movement, growth and reproduction. Air temperature, on the other hand, often varies less over the same distance. This means that there is less opportunity for ectotherms to exploit this variation through thermoregulatory behaviour, although they may hide in burrows or other types of shelters to control their body temperature in relation to air temperature. Insects can move between sunlit and shaded areas to control their body temperature and enhance their fitness by improving efficiency in foraging, escaping predators, and finding mates. However, basking in sunshine often carries costs including an increased risk of predation (Chabaud et al., 2023; Pitt, 1999), and the potential for cellular damage from ultraviolet (UV) radiation (Mazza et al., 1999, Potter and Woods, 2013).

Many studies have considered the ecological responses of ectotherms under future climate change scenarios, but much of this research has focussed on changes to average air temperature (Gardner et al., 2019) or the altered timing of extreme events (e.g., Deutsch et al., 2008; but see Kearney et al., 2009). However, shifts in air temperature have interdependence with shifts in solar radiation through variation in cloudiness. In recent decades, warming temperatures have increased the air’s moisture holding capacity, leading to greater cloud cover over much of the earth and a lowering of the diurnal temperature range (Dai et al., 1999; Cox et al., 2020). Clouds attenuate shortwave radiation, and so reduce the radiation absorbed by organisms. This can limit body temperature, even if air temperature is high (Gardner et al., 2024). The fact that temperature, radiation, and cloudiness are interrelated, and the historical focus on responses to these factors in isolation prompted our research into how temperature and radiation interact to determine life history traits.

In this study, we took advantage of altitudinal differences in air temperature and manipulated solar radiation levels through shading treatments and measured the effects on the life history traits development time and mass at maturity in an annual insect, the field cricket, *Gryllus campestris*. Growth and development rates of immature insects depend on body temperature (Kingsolver, 1989) and are major components of individual fitness. However, there is an inevitable trade-off between emergence date and adult size because insects could opt to emerge earlier after a shorter period of growth. Building a larger body must also trade-off against other fitness related traits because feeding involves exposure to risks such as predation, and resources used for growth could be used for other traits such as the immune system. In the early spring, temperate insects are expected to be well below their thermal optima and hence lower air temperature and reduced access to solar radiation are both expected to be associated with either lower adult mass or later emergence.

*G. campestris*, like many other insects, has a widespread distribution across altitudes (Panagiotopoulou et al., 2016). High altitudes are much colder than lower altitudes, which may mean that solar heat gain during the spring is even more important in crickets living at higher altitudes. A broad altitudinal range may be possible because individuals express different phenotypes in different environments (Acasus-Rivero et al., 2019), or because geographically adapted phenotypes may evolve, ie., there is local adaptation. Understanding the potential for local adaptation (Williams, 1966) allows us to be more general about responses across the species’ range. This will be important for predicting responses to climate change (Franks & Hoffmann, 2012; Schilthuizen & Kellermann, 2014) and potential future distributions. In a previous laboratory study (Tregenza et al., 2021) we investigated the growth rate of nymphs immediately after hatching comparing the offspring of parents from high and low altitudes.

We found that there was an interaction between rearing temperature and whether parents were from high or low altitude. Specifically, we found that newly hatched nymphs kept at a high temperature grew faster during the first few weeks of their lives when their parents were from low altitudes relative to those from high altitudes. However, we have no information about the growth that occurs post-diapause, nor about the independent effects of air temperature and solar radiation and possible local adaptation of responses to these factors. In this study, we tested for possible genetic components associated with the impact of environmental conditions on post-diapause growth and mass at maturity by including crickets from high and low altitude genetic lineages in all treatment groups. Our prediction is that there will be an interaction between genetic background and air temperature or solar radiation availability affecting growth rate and mass at maturity.

## Methods

### Study species

The field cricket *Gryllus campestris* is a typical annual temperate insect widely distributed in open grassland habitat from north Africa to northern Europe (Hochkirch et al., 2016). Eggs hatch from May to July and nymphs remain active until November when they enter winter diapause as late-stage instar nymphs. They emerge in early spring to resume foraging and growth and undergo one or two more nymphal moults before becoming adults from mid-April to mid-May. Both nymphs and adults use behavioural thermoregulation (sun basking) to raise their body temperature. Dark colouration may substantially reduce the basking time required to reach optimal body temperatures as observed in other insects (e.g., grasshoppers, Forsman, 2000). During our study period, there was ample sunshine availability for all cricket nymphs. Previous modelling work for *G. campestris* adults has suggested that at an air temperature of 20°C, body temperature could be up to 25°C lower under low solar radiation (∼100 W/m^2^) compared to high solar radiation (∼600 W/m^2^) (Gardner et al., 2024).

### Air temperature and shading effects on development time and adult mass

In spring 2021, we trapped crickets from ten populations in Asturias and Cantabria (Northern Spain) (Table 1).

**Table 1.** Location of cricket collection sites in 2021 in North Spain, with indications of capture date. Population is the ID of each population. Coordinates for each location are included in UTM format. Date shows the date when the crickets were collected at each site. Low altitude sites were <170m above sea level (L01-L05), high altitude sites were >1120m above sea level (L06-L10).

| Population | Location | UTM X | UTM Y | Altitude (m above sea level) | Date |
| --- | --- | --- | --- | --- | --- |
| L01 | Trelles | 683731 | 4817456 | 85 | 20-May-2021 |
| L02 | Laneo | 730627 | 4806304 | 66 | 27-Apr-2021 |
| L03 | Trubia | 277565 | 4820128 | 75 | 26-Apr-2021 |
| L04 | Hontoria | 345113 | 4812719 | 46 | 01-May-2021 |
| L05 | Bielva | 381517 | 4795578 | 119 | 05-May-2021 |
| H06 | Brañas de Abajo | 707218 | 4768328 | 1240 | 08-May-2021 |
| H07 | Valle de Lago | 729089 | 4772238 | 1300 | 07-May-2021 |
| H08 | Llanuces | 263209 | 4782845 | 1118 | 05-May-2021 |
| H09 | Peña Mayor | 296857 | 4794393 | 1126 | 06-May-2021 |
| H10 | Tarna | 319128 | 4774872 | 1121 | 09-May-2021 |

We bred these crickets in the lab, and between 26/6/2021 and the 20/7/2021 we released small nymphs into 120 outdoor translucent polypropylene boxes with an area of 0.2m^2^, a depth of 28cm, and a 14cm layer of soil with natural grass growing on it. To manipulate ambient air temperature, boxes from each source population were divided equally between a site at 70m above sea level (60 boxes) and a site at 1300m above sea level, 58km to the southwest (60 boxes). Ambient air temperature was ∼4°C lower at the higher altitude site compared to the lower altitude site (daily mean temperature at high altitude site during the period of our study: 9.6°C, daily mean temperature at low altitude site: 13.4°C, t test, t=-9.86, df=705, p=<0.001 (supplementary material, Figure S1). Solar radiation can change with altitude (Buckley et al., 2013), but between our sites, solar radiation was very similar during the period of our study (daily mean radiation at high altitude site: 140W/m^2^, daily mean radiation at low altitude site: 134 W/m^2^, t test, t=1.05, df=707, p=0.29). We hereafter refer to the high and low altitude sites as the low and high air temperature sites, respectively.

Boxes were initially covered with 0.5mm steel mesh lids, as small nymphs can climb up the inside of the box walls and escape. In mid-October, when nymphs were larger and became unable to climb up the plastic walls, we replaced the thin mesh lids with wider 12mm mesh lids. These lids prevented predation by birds but did not cast any appreciable shade (supplementary material, Figure S2). To manipulate solar radiation, we shaded half of the boxes with a layer of plastic mesh which had a flat profile with 1mm thick strands, forming rectangular holes of 4mm x 5mm inner size. We used a pyrometer (Davis instruments, Vantage Pro 2 (www.davisinstruments.com)) to determine that this mesh reduced the solar radiation hitting the soil inside the box to about 60% of the value in the unshaded boxes (supplementary material, Figure S2). This is similar to the solar radiative reduction of stratus clouds (∼40-80%).

From early April 2022, before adults began to emerge, we started visiting both rearing sites at intervals of a maximum of 11 days (range: 1-11 days, mean: 4 days) to record the date of emergence of new adults and to remove and weigh them. Any cricket with signs of very recent emergence (still white or seen moulting by the day of observation) was assigned that date as its emergence date and others were assigned the mid-date between the previous visit and when we found them as adults. Some individuals were very difficult to see or to catch during our visits. For those, we either assigned an emergence date as the midpoint within the period between the day when we saw the first adult at that rearing site, and the date when we saw it for the first time, or we did not assign any emergence date. Adults removed from the rearing boxes were weighed within the following 2 days with a 0.01 g accuracy balance.

Please see supplementary material for further details of all methods.

### Analysis

We compared the development time (number of days from 1st January to the date of emergence as an adult) and initial adult mass (in grams) between shaded and unshaded boxes at high and low air temperature sites using linear models. Site (air temperature) and shading treatment and their interaction, and altitude of origin were included as predictors, and sex was included as an additional predictor in the adult mass model as females are on average larger than males.

Diagnostic plots were checked and showed that model assumptions of normality and homoscedasticity were met. A potential relationship between size at adulthood and emergence date, which has been reported previously (Ritz and Köhler, 2007) was examined using a Kendall’s rank correlation test.

All statistical analyses were carried out in R, version 4.3.1 (R Core Team, 2023).

## Results

Development time (Figure 1a) was strongly affected by air temperature: crickets at the high air temperature site emerged on average 10 days earlier than those in the low air temperature site (t_181_=6.94, p<0.0001). Development time was not significantly affected by whether or not crickets were in shade (est=-1.46 days, t_181_=0.94, p=0.35). Altitude of origin did not affect development time (t_181_=0.70, p=0.49). Please see Table S1 in the supplementary material for all statistical results.

**Figure 1.**
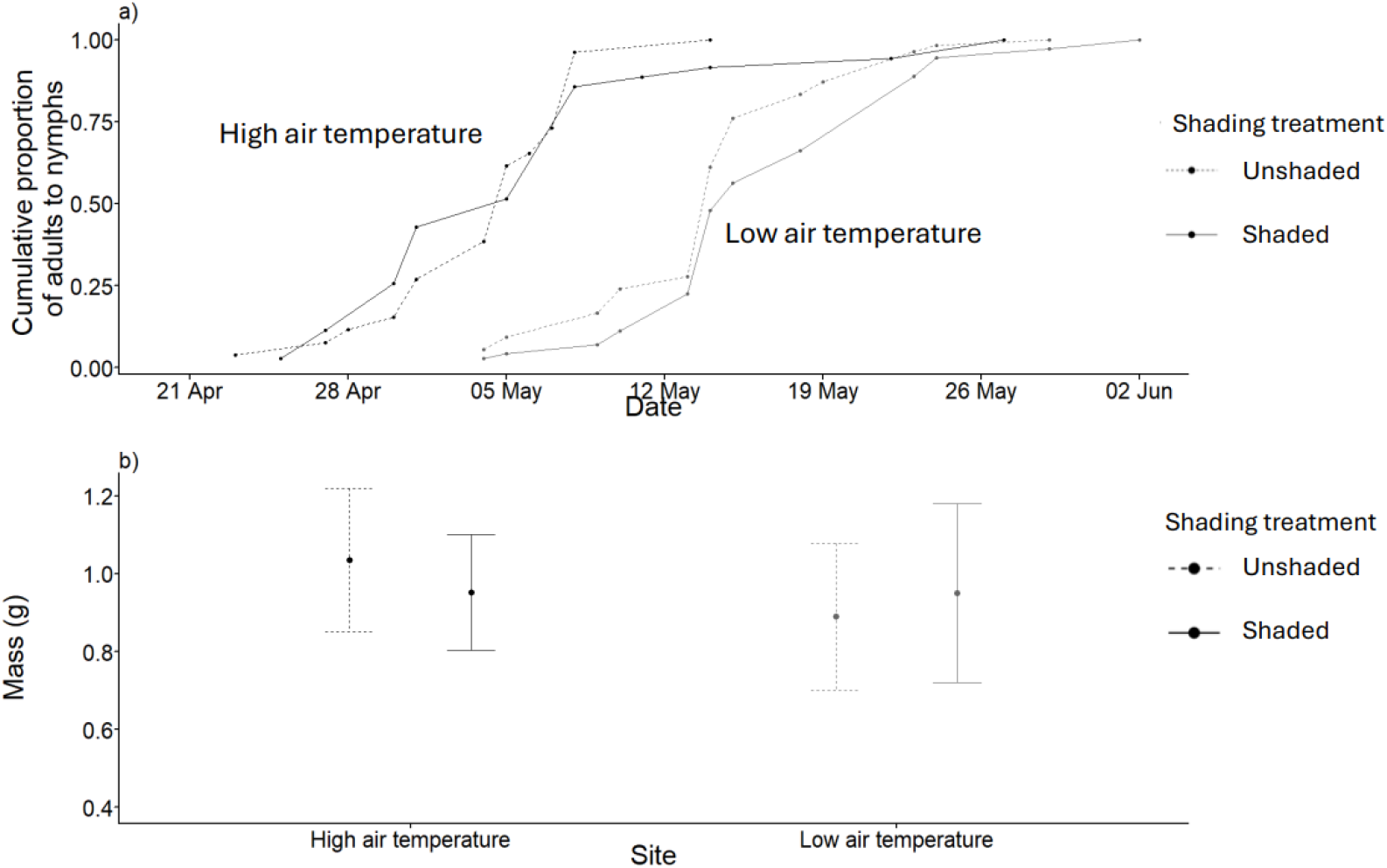
a) cricket emergence date (cumulative proportion of all adults to nymphs) and b) adult mass (in grams) for high (black) and low (dark grey) air temperature sites for unshaded (dashed lines) and shaded (solid lines) treatments. In b), points show mean values, whiskers extend to one standard deviation above and below the mean value. Sample sizes as follows: high air temperature site: shaded=35, unshaded=26; low air temperature site: shaded=71, unshaded =54.

Adult mass was lower at the low air temperature site (Figure 1b) (est=-0.25g, t_177_=-3.81, p<0.001). Adult mass was also dependent upon an interaction between the air temperature and the availability of solar radiation: at low air temperature, shaded crickets had greater mass than unshaded crickets whereas at high air temperature, shaded crickets had lower mass than unshaded crickets (t_177_=2.29, p=0.02). Males had lower mass than females (mean mass of females: 0.97g, mean mass of males: 0.92g, est=-0.19g, t_177_=-2.46, p=0.156). Altitude of origin did not affect adult mass (t_177_=-0.88, p=0.38). Please see Table S2 in the supplementary material for all statistical results.

We found no significant correlation between adult mass and emergence date (τ=-0.06, z=-1.11, p=0.27).

## Discussion

### Effects of air temperature

Nymphs became adult substantially later at lower air temperatures. Development time is an important life history trait and is expected to be under strong selection. Earlier adult maturity has the potential to reduce generation time and reduces the period during which individuals are exposed to predators and other mortality risks prior to reproductive maturity. Hence earlier reproductive maturity tends to increase the probability that an individual passes on its genes before it dies. However, in annual species the fixed generation time means that benefits of earlier maturity for parents are weighed against the requirement for their first offspring to then live longer to remain in sync with the annual cycle. Additionally, seasonal effects create an optimal period for egg laying and nymphal development which means that very early or late adult maturity is likely to be selected against, and so overall, we would expect stabilising selection on emergence date. We previously identified a positive relationship between early emergence and lifespan in *G. campestris* males (Rodríguez-Muñoz et al., 2019). However, a subsequent analysis measuring selection as the number of adult offspring produced in the following year (Rodríguez-Muñoz et al., submitted) reveals that across 8 years, selection on emergence date tends to be neutral.

Another study (Ritz and Köhler, 2007) reported earlier emergence in larger nymphs in a wild population of *G. campestris*. However, this assertion was based on a post-hoc analysis showing a decline in the mean size of crickets over a 10-day period at the beginning of the breeding season. The approach of observing a trend over part of a time series and analysing that trend in isolation from the subsequent increase in average size carries a high risk of a Type 1 error. Our analysis, in which we can identify specific individuals and measure the correlation between body size and emergence date fails to find such a relationship. There is inevitably a trade-off between development time and size at maturity, but whether this leads to a particular relationship between these traits depends on costs and benefits relating to the two traits (e.g. Gershman et al., 2022).

In many ectotherms, individuals grow more slowly but are larger as adults in colder environments (Atkinson (1994, 1995)), an effect known as the temperature-size rule (Atkinson 1996). However, in this annual species, we found that despite developing more slowly, nymphs matured at a slightly lower adult mass in the lower air temperature treatment. This may be because temperatures at the high air temperature site were not high enough to compromise growth efficiency, or more generally, because of the complex relationship between temperature and body size (Angiletta and Dunham, 2003). An earlier analysis of a single year at our low altitude site found that larger *G. campestris* adults had significantly higher lifetime reproductive success (Rodríguez-Muñoz et al., 2010). However, subsequent analysis across 8 years indicates that this is not a general pattern, and that there is no consistent directional selection in favour of larger body size (Rodríguez-Muñoz et al., submitted). It is possible that at higher altitudes, there is a trade-off whereby lower mass becomes more advantageous (Olalla-Tárraga and Rodríguez, 2007) since heating rates are determined by body mass (as body mass decreases, body surface area gets proportionally larger, which contributes to increased rates of heat gain, which would be beneficial in a colder environment). However, this effect requires further investigation. We do note that the difference in mass between the high and low ambient air temperature site, although significant, was relatively small compared to the very large effect on development time, so it does appear that crickets are optimising mass at adulthood at the expense of taking longer to develop

### Effects of solar radiation

Our shading treatment imposed a significant reduction in solar radiation on our crickets, so we were expecting a large effect on the key life history traits of development rate and mass at adulthood. However, crickets developed at similar rates in the shaded and unshaded treatments, suggesting that they can compensate for suboptimal radiative environments. It is likely that shaded crickets compensated for reduced solar radiation by increasing the amount of time they spent sun basking (Gardner et al., in press). This assertion is supported by the observation that *G. campestris* adults spend more time in the sun when under partial shade (Gardner et al., in press); if nymphs behave in the same way, then this would drive the patterns we observe.

Assuming that our nymphs are indeed compensating for partial shade by extending their basking time, this capacity to compensate means that unshaded crickets at high altitude are forgoing the opportunity to bask more, i.e. there is sun available that they are not using. Predation is a likely driver of basking avoidance (Huey and Slatkin, 1976), particularly early in the season when the sward height is very short and so predation risk from birds is likely to be high. Pitt (1999) for example, found that grasshoppers utilise different microhabitats (positions in vegetation) to balance the trade-off between reducing mortality from predators and experiencing greater food availability, and warmer conditions. Several studies have also found that solar UVB radiation can negatively impact development, survival, and reproduction in insects (Mazza et al., 1999; Potter and Woods, 2013). When shaded, the benefits of basking (i.e., to maintain development time) seemingly outweigh these potential costs. In our study natural predation by birds and mammals was prevented because the boxes containing the crickets were covered with a thin mesh. This meant that increased basking by shaded crickets was not penalised by potentially higher predation rates, so if there are interactions between shading, basking behaviour and predation mortality (e.g., Battisti et al., 2013) we would not expect to observe it directly in our study.

Shading had a negative effect on adult mass at the high air temperature site, but a positive effect on adult mass at the low air temperature site, although the actual difference in adult mass between treatments was small. Battisti et al. (2013) report a similar effect of shading on first-instar colonies of *Thaumetopoea pinivora*; shading did not affect development rate, but negatively impacted growth rate. The authors suggest that the greater size achieved for the same development time in unshaded colonies could increase their capacity to resist predation. Predation risk does not seem to be related to size in field crickets – the majority of predation events that we observe are by large predators such as robins and shrews (Rodríguez-Muñoz et al., 2011) which can handle the largest crickets. However, it may be that in response to the shading treatment, cricket nymphs at the low air temperature site accrued benefits from increased low-level growth rates that resulted in larger body mass because being larger in cold habitats can increase organisms’ ability to conserve heat (Meiri, 2011). Equally, they may have suffered by being larger in a colder location with compromised basking opportunities due to reduced rates of heat gain at higher mass.

### Local adaptation

We found no evidence for local adaptation influencing responses to air temperature or shading in post-diapause growth. This suggests that crickets from high altitude genetic populations are not genetically adapted to express different behavioural thermoregulation strategies in anticipation of cooler conditions (for instance basking more). This finding contrasts with our earlier observation (Tregenza et al. 2021) of an interaction between genetic background and rearing temperature affecting the growth rate of very small nymphs in the period immediately post-hatching. Our earlier observation suggests that local genetic adaptation to altitude can occur in the populations that we studied. Therefore, the lack of any evidence of such adaptation in this study indicates that there is not very strong selection for differences in growth rate, mass at adulthood or interactions between these traits and air temperature and solar radiation availability in post-diapause nymphs. This finding is somewhat unexpected because the much shorter breeding seasons at high altitude might be expected to favour more rapid growth when heat is available. However, precise predictions are hard to make because a shorter breeding season also means that there is a longer pre-adult phase which provides more time for growth before mates are available.

### Wider implications

Differences between altitudes are not confined solely to air temperature. For example, oxygen concentration is lower at higher altitude which may affect respiration, and lower air density may reduce convective heat loss of insects (Dillon et al., 2006). However, these differences are only likely to be significant at altitudes higher than those we were using; convective heat loss at 70 and 1300m above sea level is almost identical at the low wind speeds our crickets were exposed to (see Figure 3 in Dillon et al., 2006). Therefore, we believe air temperature likely is the most important factor influencing life history traits between altitudes in this study.

Although we report effects of ambient temperature and shading on life history traits in nymphs, how these factors affect lifetime fitness (in term of the probability that an individual of a given phenotype will contribute to subsequent generations (Lailvaux and Husak, 2014)), remains unknown. It would be extremely difficult to track individual crickets from nymphs to adults in a natural context. However, in future studies we could analyse how air temperature and access to solar radiation affect fitness or fitness-related traits in adult crickets. There are likely complex trade-offs and strategies to optimise fitness in a particular environment (Lahondère, 2023).

Better understanding of the possible fitness consequences of thermal environments for ectotherms, including possible trade-offs of increased basking activity under different temperature regimes (e.g., Ma et al., 2018), will help to uncover the drivers governing their distributions (Larsson, 2002). This will benefit numerous fields, including ecology and biogeography, with applications in the assessment of pest risk (e.g., Mezei et al., 2019) and the possible consequences of climate change. Given that temperature and solar radiation (through changes to cloud cover) are important components of climate change (although future trends in cloudiness and radiation are complex to predict) (IPCC, 2013, Cox et al., 2020), understanding how warmer temperatures and altered basking opportunities (due to changes in cloud cover) might interact and impact ectotherms becomes crucial. The opportunity to bask is considered an important determinant of the spatial distribution of ectotherms (Battisti et al., 2013). Thus, behavioural thermoregulation is very likely to be an important mechanism to ‘buffer’ ectotherms against negative impacts of climate change (Kearney et al., 2009). Our results suggest that developing crickets forgo some opportunities to gain energy from the sun, because if shaded they can compensate to maintain development rate. Our observation that the direction of the effect of shade on adult mass depends on the air temperature means that changes in cloud cover may impact insects differently in warmer and cooler areas.

## Ethics statement

This study has been approved by the University of Exeter’s Research Ethics Panel, approval number 513752. All crickets used in the study will live out their natural lives in the wild.

## Data accessibility

The data used in the analysis are deposited in Zenodo: https://doi.org/10.5281/zenodo.11090808 and will be made publicly accessible on manuscript acceptance**.

**Zenodo link for peer review: https://tinyurl.com/y4v89x75

## Acknowledgments

Thanks to Emma Álvarez Alba for generously providing us with land for the high-altitude rearing site and for contributing to maintenance of the crickets. Thanks to Paul Hopwood for comments on an earlier draft of the manuscript. This work was supported by the Natural Environment Research Council (NERC); standard grant: NE/V000772/1.

## Funding

This work was supported by Natural Environment Research Council grants NE/R000328/1 and NE/V000772/1.

## Competing interests

The authors declare no competing interests.

## Notes

https://tinyurl.com/y4v89x75

### Competing Interest Statement

The authors have declared no competing interest.

### Summary of Updates

I have added author Alfredo F Ojanguren who should have been listed on the original version in recognition of his contribution to the manuscript. He has contributed to the work reported in the paper and consented to its submission.

