## Supplementary Material for "How air temperature and solar radiation impact life history traits in a wild insect"

**Collecting sites**

During the spring 2021, we trapped adult crickets or last instar nymphs from ten different populations in Asturias and Cantabria (North Spain). Five of the populations were located at altitudes below 170m and five above 1120m (see Table 1 in the main text). On the day of capture, we stored the crickets in 20l boxes with egg cardboard for shelter, food (standard rodent diet) and water (a 5ml glass vial filled with water and closed with a cotton plug) and moved them to the lab within the following 2 days.

**Cricket rearing and breeding**

At the laboratory, we isolated up to ten males and ten females per population (depending on the numbers available) in 2l plastic boxes, prioritising adults over nymphs. Each box had a window covered with a thin metal mesh for ventilation, a piece of egg cardboard for shelter and food and water ad libitum. We stored the excess females (for those populations with more than 10 available) in groups of 4-5 using the same type of 2l boxes, whereas males in excess remained grouped in the 20l boxes used during capture. We placed the boxes in a shelving within an open outdoor shed, where they remained in the shade, with ambient light and temperature. The only exception to this rearing protocol was population H07, for which we only captured nymphs. To promote adult emergence under natural conditions, we placed these nymphs in open-top plastic boxes located outdoors. The boxes had a 14cm layer of soil with natural grass growing on it, and an area of $0.2m^2$. We used three boxes per sex, with 2-3 females or males in each box, and fed them with sunflower seeds. Once they emerged as adults, they were isolated following the protocol described above for the rest of the populations. Males and females maintained in groups were only included in the protocol of isolation and mating (see below), whenever we had to replace any of the originally isolated individuals because of death or failure to mate and/or produce eggs.

Crickets collected as adults might have mated in the wild before we trapped them and so would be able to lay fertile eggs straightaway. However, because we caught them at the start of the breeding season, it is likely that many of the females were still virgin. To make sure that the experimental females had mated with at least one male, and so were able to lay fertile eggs, from mid-May we carried out mating trials with all the isolated females. At each mating trial, we put one pair of crickets from the same population into a 0.5l plastic cup, with a piece of paper on the bottom to provide traction. We left the pair together in the cup and checked every few minutes or so over a period of 1-3 hours, to see if the female had a spermatophore attached under her ovipositor. After their first mating trial, we weighed the females, took a picture for thorax measurement, and moved them to a room indoors (using the same rearing box) with natural light and 25°air temperature, recording which females had mated. In nature, females lay eggs in the ground using their long ovipositor. In the lab, we provide them with a small Petri dish (3 cm in diameter and 1 cm in depth) filled with wet sand. For females that did not mate on any given mating trial, we carried out further mating trials (maximum 3-5 trials) on subsequent days. After those trials, we discarded any female that did not mate in the lab and did not lay any eggs.

Every 1-3 days we sieved the sand of each female's dish to collect any eggs and replaced it with a new dish with clean sand. We distributed the eggs of each sieved dish on a wet cotton-wool pad placed in a Petri dish for incubation, trying to avoid clumps of eggs that could favour fungal growth. We left the dishes in the same room used to rear the laying females. After 4-5 days, we checked incubation dishes daily (at 25°C incubation takes around 9 days) to remove hatchlings and transfer them to a plastic box like those used for the isolated females. We reared all the hatchlings from consecutive laying events in the same box (i.e. only one hatchling box per female), and in the same room as laying females and incubation dishes.

After 2-3 weeks growing in the boxes, we released a sample of about 50 small nymphs per female into each of four boxes outdoors (the same boxes used to rear the H07 nymphs until adulthood), two of them located in our WildCrickets meadow at 70 m above sea level, and two in a meadow at 1300 m above sea level 58km to the southwest of the WildCrickets meadow. Solar radiation is similar between sites but the air temperature is lower at higher altitude (figure S2). For a small number of females that did not produce 200 nymphs, we divided all the available nymphs into four groups and released each group into one of the boxes. Each box hosted nymphs from a single female. Initial box density was 25-32 nymphs in 110 boxes, and 9-23 in 10. Releasing took place in two batches, on the $26^{th}$ of June and the $20^{th}$ of July, for the earlier and later hatched nymphs, respectively.

On the day of releasing the nymphs in the boxes outdoors, we covered all boxes with thin metal 0.5mm mesh lids to prevent small nymphs from escaping. In mid-October, we replaced the thin mesh lids with wider 12mm mesh lids. By that time, nymphs were unable to climb up the slippery walls of the boxes; the lids prevented predation attacks from birds but only caused only very small degree of shade (Figure S1).

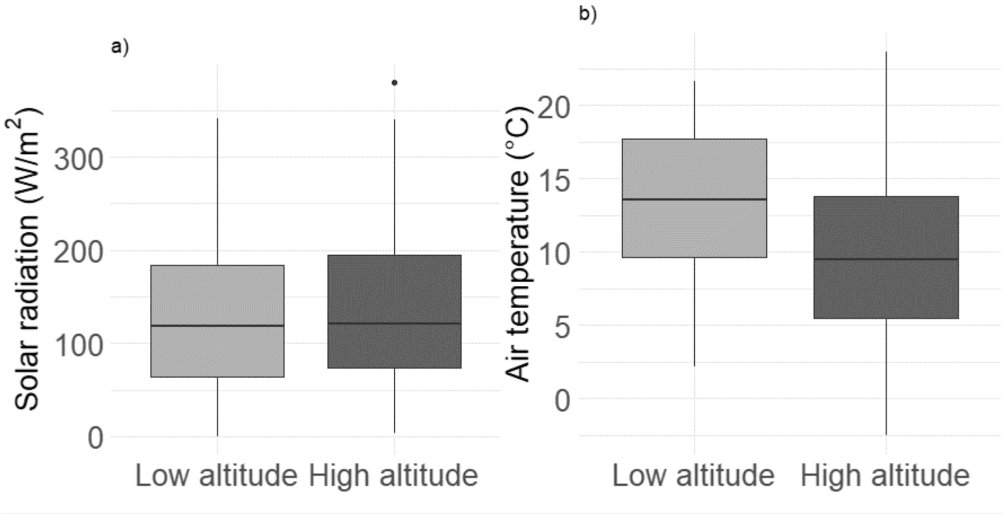


**Figure S1:** Summary of a) mean daily radiation ($W/m^2$) and b) mean daily air temperature (°C) at the low and high altitude site. Data are for 01/06/2021-31/05/2022, corresponding to the period from when crickets went into boxes up until the median emergence date as an adult (see methods in main text). Data were recorded at the low and high altitude site using Davis instruments, Vantage Pro 2 weather stations (www.davisinstruments.com). Solid lines crossing the boxes are the median value. The lower and upper hinges of each box correspond to the first and third quartiles (the 25th and 75th percentiles). Respectively, the upper and lower whiskers extend to the largest and smallest values no more than 1.5 times the interquartile range. Individual points are outliers.

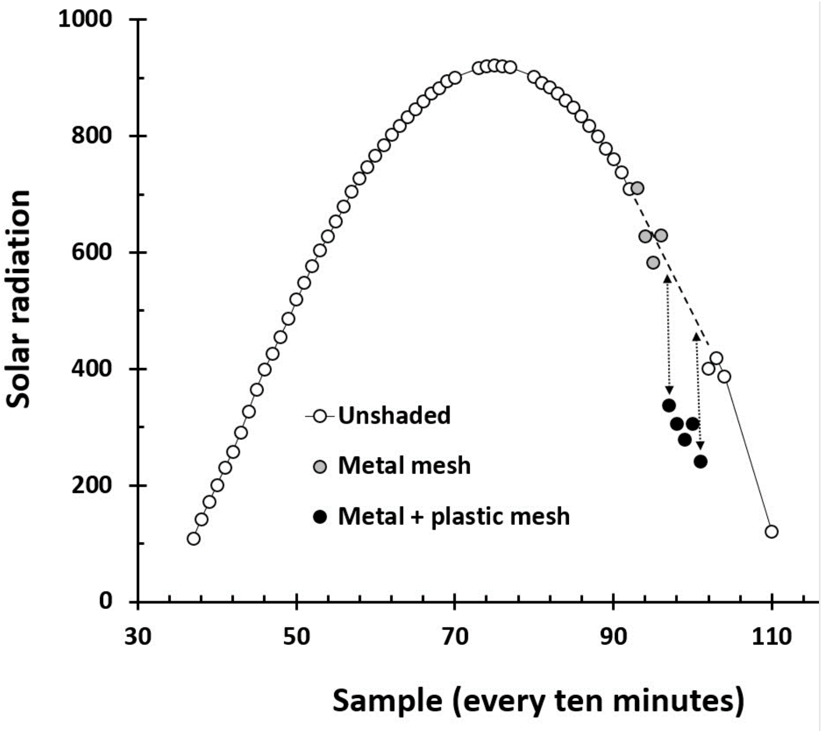


**Figure S2: Solar radiation during a day of full sun measured at 1.5 m above the ground using Davis instruments, Vantage Pro2 Plus pyrometer**. Grey dots show when a metal mesh was placed above the pyrometer which has a minimal impact on solar radiation. Black dots show a period where the plastic mesh treatment was applied; solar radiation can be seen to reduce by ~40% relative the expected value when unshaded.

**Table S1: Results of a Gaussian linear model analysing predictors of development time** (calculated as the number of days after Jan 1[st] that a cricket emerged as adult)**.** Total number of observations: 186.

| **Variable** | **Estimate** | **Std. error** | **P value** |
|---|---|---|---|
| Intercept | 122.81 | 1.47 | <0.0001 |
| Site (low air temperature) | 10.04 | 1.45 | <0.0001 |
| Treatment (shaded) | 1.46 | 0.94 | 0.35 |
| Altitude of origin (<170m) | 0.69 | 0.70 | 0.49 |
| Site (low air temperature) * treatment (shaded) | 1.50 | 0.79 | 0.43 |

**Table S2: Results of a Gaussian linear model analysing predictors of adult mass (in grams).** Total number of observations: 186.

| **Variable** | **Estimate** | **Std. error** | **P value** |
|---|---|---|---|
| Intercept | 1.16 | 0.06 | <0.0001 |
| Treatment (shaded) | -0.13 | 0.08 | 0.10 |
| Site (low air temperature) | -0.26 | 0.07 | <0.0001 |
| Sex (male) | -0.19 | 0.08 | 0.02 |
| Altitude of origin (<170m) | -0.03 | 0.03 | 0.38 |
| Treatment (shaded) * site (low air temperature) | 0.21 | 0.09 | 0.02 |
| Treatment (shaded) * sex (male) | 0.09 | 0.10 | 0.39 |
| Site (low air temperature) * sex (male) | 0.20 | 0.09 | 0.04 |
| Treatment (shaded) * site (low air temperature) * sex (male) | -0.14 | 0.12 | 0.26 |
